# Perturbation of effector and regulatory T cell subsets in Myalgic Encephalomyelitis/Chronic Fatigue Syndrome (ME/CFS)

**DOI:** 10.1101/2019.12.23.887505

**Authors:** Ece Karhan, Courtney L Gunter, Vida Ravanmehr, Meghan Horne, Lina Kozhaya, Stephanie Renzullo, Lindsey Placek, Joshy George, Peter N Robinson, Suzanne D Vernon, Lucinda Bateman, Derya Unutmaz

## Abstract

Myalgic encephalomyelitis/chronic fatigue syndrome (ME/CFS) is a debilitating disorder of unknown etiology, and diagnosis of the disease is largely based on clinical symptoms. We hypothesized that immunological disruption is the major driver of this disease and analyzed a large cohort of ME/CFS patient or control blood samples for differences in T cell subset frequencies and functions. We found that the ratio of CD4+ to CD8+ T cells and the proportion of CD8+ effector memory T cells were increased, whereas NK cells were reduced in ME/CFS patients younger than 50 years old compared to a healthy control group. Remarkably, major differences were observed in Th1, Th2, Th17 and mucosal-associated invariant T (MAIT) T cell subset functions across all ages of patients compared to healthy subjects. While CCR6+ Th17 cells in ME/CFS secreted less IL-17 compared to controls, their overall frequency was higher. Similarly, MAIT cells from patients secreted lower IFNγ, GranzymeA and IL-17 upon activation. Together, these findings suggest chronic stimulation of these T cell populations in ME/CFS patients. In contrast, the frequency of regulatory T cells (Tregs), which control excessive immune activation, was higher in ME/CFS patients. Finally, using a machine learning algorithm called random forest, we determined that the set of T cell parameters analyzed could identify more than 90% of the subjects in the ME/CFS cohort as patients (93% true positive rate or sensitivity). In conclusion, these multiple and major perturbations or dysfunctions in T cell subsets in ME/CFS patients suggest potential chronic infections or microbiome dysbiosis. These findings also have implications for development of ME/CFS specific immune biomarkers and reveal potential targets for novel therapeutic interventions.

## Introduction

Myalgic encephalomyelitis/chronic fatigue syndrome (ME/CFS) is a highly debilitating illness often characterized by symptoms such as post-exertional malaise or severe fatigue not alleviated by rest, muscle and joint pain, sleep problems, hypersensitivity to sensory stimuli, and gastrointestinal symptoms (Holgate et al., 2011;Yancey and Thomas, 2012;Bested and Marshall, 2015). ME/CFS is thought to afflict up to two million individuals in the US alone, with severe long-term disability and negative impacts on quality of life (Valdez et al., 2018). The specific cause and biological basis of ME/CFS remain elusive. Lack of understanding of biological pathways leading to this syndrome is also a major impediment in developing specific therapies and reliable biomarker-based diagnostic tests (Klimas et al., 2012).

While the causes of ME/CFS are likely to be multifactorial, a common history of initial infectious agents, including viral (e.g. EBV) and bacterial (e.g. Lyme Disease) infections, have been associated with triggering the disease (Hickie et al., 2006;Katz et al., 2009). Indeed, mounting evidence in ME/CFS patients implicates a significant role for immunological abnormalities that are thought to contribute to disease progression and/or maintenance of the chronic symptomatic state (Fletcher et al., 2010;Brenu et al., 2012;Curriu et al., 2013;Brenu et al., 2014;Mensah et al., 2017). Studies of the immune system of ME/CFS subjects have revealed many abnormalities, including disruptions in the number and function of T cell subsets, B cell and natural killer (NK) cells (Fletcher et al., 2010;Brenu et al., 2012;Curriu et al., 2013;Brenu et al., 2014); changes in T-cell or innate cell cytokine secretion (Torres-Harding et al., 2008;Broderick et al., 2010;Bansal et al., 2012); changes in humoral immunity (Prinsen et al., 2012) and inflammatory immune signaling (Aspler et al., 2008); and higher frequencies of various autoantibodies (Ortega-Hernandez and Shoenfeld, 2009). In particular, T cells are responsible for orchestrating and modulating an optimal immune response, either through their effector or regulatory functions. Thus, perturbations in T cell subsets or in effector or regulatory functions during ME/CFS (Lorusso et al., 2009;Rivas et al., 2018) can result in an immune disruption or unwanted immune responses.

Here we found perturbations in CD8+ T cells, NK cells, Treg cell frequencies and effector functions of Th17 and MAIT cells in ME/CFS patients. Some of the immune parameter differences, such as CD8+ effector memory T cells and NK cells, were relatively different in younger patients, but not in patients older than 50 years, when compared to healthy controls. In addition, using significant immune features and a machine learning algorithm, we could identify ME/CFS patients among healthy controls with very high sensitivity and specificity. Together, our findings reveal multiple chronic T cell dysfunctions in ME/CFS, suggesting a link to chronic infections or a disruption of the microbiota.

## Materials and Methods

### Participants

All subjects were recruited at Bateman Horne Center, Salt Lake City, UT, based on were who met the 1994 CDC Fukuda (Fukuda et al., 1994) and/or Canadian consensus criteria for ME/CFS (Carruthers, 2007). Healthy controls were frequency-matched to cases on age, sex, race/ethnicity, geographic/clinical site and season of sampling. Patients or controls taking antibiotics or had any infections in the prior months; taking any immunomodulatory medications were excluded from the study. The study was approved by Western IRB (Protocol number 20151965) and written informed consent and verbal assent when appropriate were obtained from all participants in this study. We enrolled a total of 198 ME/CFS patients and 91 healthy controls. Subject characteristics are shown in supplementary Table 1.

### PBMC isolation and preservation

Healthy and patient blood samples are obtained from Bateman Horne Center, Salt Lake City, UT and approved by Western IRB. Heparinized blood samples were shipped overnight at room temperature. Peripheral blood mononuclear cells (PBMC) were then isolated using Ficoll-paque plus (GE Health care) and cryopreserved in liquid nitrogen.

### Surface and intracellular staining and flow cytometry analysis

After thawing, PBMCs were counted and divided into 2 parts, 1 part for day 0 surface staining, and the other part cultured in complete RPMI 1640 medium (RPMI plus 10% Fetal Bovine Serum (FBS, Atlanta Biologicals) and 1% penicillin/streptomycin (Corning Cellgro) supplemented with IL-2+IL15 (20ng/ml) for Treg subsets day 1 surface and transcription factors staining, IL-7 (20ng/ml) for day 1 and day 6 intracellular cytokine staining and a combination of cytokines (20ng/ml IL-12, 20ng/ml IL-15, and 40ng/ml IL-18) for day 1 intracellular cytokine staining (IL-12 from R&D, IL-7 and IL-15 from Biolegend).

Surface staining was performed in staining buffer containing PBS + 2% FBS for 30 minutes at 4°C. When staining for chemokine receptors the incubation was done at room temperature. Antibodies used in the surface staining are CD3 (UCHT1 clone, Alexa Fluor 532, eBioscience), CD4 (OKT4 clone, Brilliant Violet 510), CD8 (RPA-T8 clone, Pacific Blue or Brilliant Violet 570), CD19 (HIB19 clone, Brilliant Violet 510), CD45RO (UCHL1 clone, Brilliant Violet 711, APC/Cy7, or Brilliant Violet 570), CCR7 (G043H7 clone, Alexa Fluor 488), 2B4 (C1.7 clone, PerCP/Cy5.5), CD14 (HCD14 clone, Alexa Fluor 700), CD27 (O323 clone, PE/Cy7, Brilliant Violet 605, or Alexa Fluor 647), CCR6 (G034E3 clone, Brilliant Violet 605), CD161 (HP-3G10, Brilliant Violet 421), Vα7.2 (3C10 clone, PE) (all from Biolegend).

For intracellular cytokine staining, cells were stimulated with PMA (40ng/ml for overnight cultured cells and 20ng/ml for 6 days cultured cells) and Ionomycin (500ng/ml) (both from Sigma-Aldrich) in the presence of GolgiStop (BD Biosciences) for 4 hours at 37^°^C. For cytokine secretion after stimulation with IL-12+IL-15+IL-18+, GolgiStop was added to the culture on day 1 for 4 hours. Stimulated or unstimulated cells were collected, stained with surface markers including CD3, CD4, CD8, CD161, Vα7.2, CD45RO, CCR6, and CD27 (Biolegend) followed by one wash with PBS (Phosphate buffer Saline) and staining with fixable viability dye (eBioscience). After surface staining, cells were fixed and permeabilized using fixation/permeabilization buffers (eBioscience) according to the manufacturer’s instruction. Permeabilized cells were then stained for intracellular IFNγ (4S.B3 clone, APC/Cy7), TNFα (Mab11 clone, PE/Dazze 594), GranzymeA (CB9 clone, Alexa Fluor 647, Alexa Fluor 488), IL-17A (BL168 clone, Alexa Fluor 488, Brilliant Violet 421), Foxp3 (259D clone, PE), and Helios (22F6 clone, Alexa Fluor 488) (all from Biolegend). Flow cytometry analysis was performed on SP6800 spectral cell analyzer (Sony Biotechnology) and analyzed using FlowJo (Tree Star).

### Machine learning and statistical analysis

All statistical analyses were performed using GraphPad Prism V8 software. Continuous variable datasets were analyzed by Mann-Whitney U test for non-parametric datasets when comparing clinical groups, and exact P values are reported. Spearman p was used to determine the relationship existing between two sets of data between non-parametric datasets.

The algorithms for identifying significantly different features and the RF classifier were implemented in Python 3.6.8 using Jupyter Notebook 5.0.0. The RF classifier was performed with different numbers of features of *k* = 65, 40, 10. A training set with 231 samples (80% of total samples) was selected and the remaining data corresponding to 58 samples (20% of total samples) was left as the test set. Missing values in the training and test sets were replaced by corresponding median value in the training set. The RF classifier was implemented using a 3-fold (stratified) cross validation and was trained using all 65 immune profile features, the 40 significantly different features, the top 10 significantly different features and the top 10 features among the 40 significantly different features that received the highest importance score.

There are several metrics to evaluate the performance of a classifier. Sensitivity represents the proportion of patients who were correctly identified as patients and specificity represents the proportion of healthy controls who were correctly identified as healthy. If patients are denoted by “positives” and healthy controls by “negatives”, then sensitivity and specificity are calculated as:

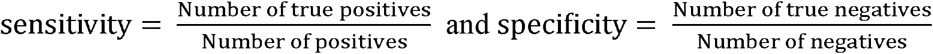

where “true positives” refer to patients who were correctly identified as patients and “true negatives” refer to healthy controls who were correctly identified as healthy.

Accuracy is a metric which shows the fraction of predictions that our classifier predicted correctly. Accuracy is calculated in terms of true positives and true negatives as:

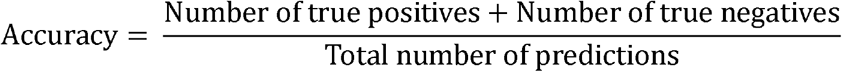

Positive (negative) predictive values are the proportion of positives (negatives) that are correctly identified as positives (negatives which are calculated as:

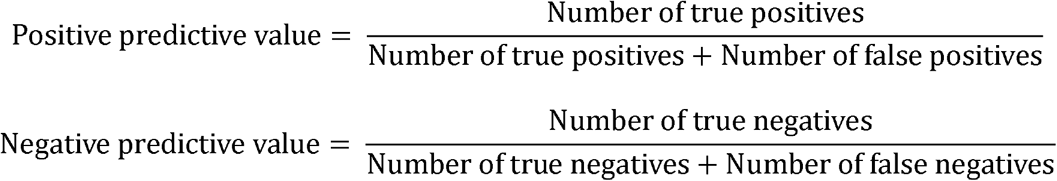

The F_l_ score measures the accuracy of test by calculating the (harmonic) mean of the sensitivity (recall) and positive predictive value (precision). The F_l_ score is defined as:

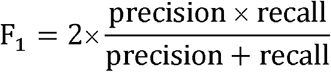

## Results

### Changes in T cells subsets in ME/CFS patient blood

To determine phenotypic and functional changes in immune cell subsets from ME/CFS patients, we developed several staining panels for flow cytometry analysis and performed immune profiling of 198 ME/CFS patients and 91 age- and sex-matched healthy controls (Supplemental Table 1). We first analyzed the main immune subsets in peripheral blood mononuclear cells (PBMCs), namely T cells, B cells, NK cells, and monocytes (Figure 1a, 1b). There was no significant difference in the percentage of overall monocytes (p=0.9), B cells (p=0.9) or T cells (p=0.1) (Fig. 1a, 1b), but the frequency of NK cells within lymphocytes was greatly reduced (p=0.0005) in ME/CFS compared to healthy controls (Fig. 1b). Within T cells, we observed that CD4+ T cell frequency was higher (p=0.0193) and correspondingly, CD8+ T cells were lower (p=0.0052) in ME/CFS subjects, and that this was reflected as a higher CD4 to CD8 ratio (p=0.0078) in patients (Fig. 1c). There was no difference in CD4-CD8-(double negative; DN) T cells (p=0.9) between controls and ME/CFS patients (data not shown).

**Figure 1.**
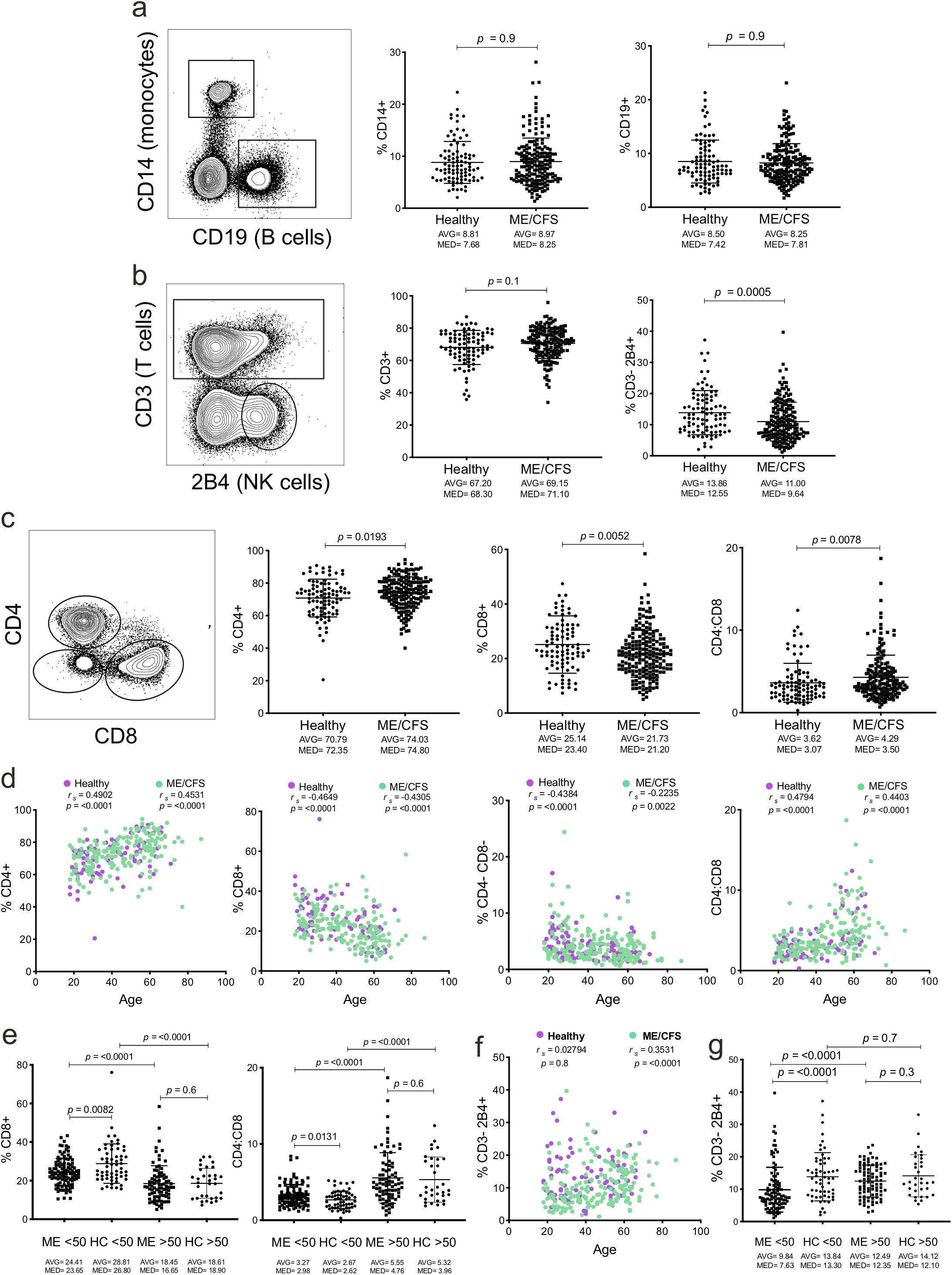
Frequency of CD4+ and CD8+ T cell subsets in ME/CFS patients and healthy controls. Analysis of frequency of main immune subsets in PBMC by flow cytometry was performed after staining and gating on (a) CD14+ (Monocytes) and CD19+ (B cells), and (b) CD3+ (T cells) and CD3-2B4+ (NK cells), and proportion of each subset frequency as a portion of PBMC are shown for each subject (right panels). (c) Frequencies of CD4+ and CD8+ T cells and their ratio were analyzed within CD3+ T cell gates. (d) Correlation of T cell subsets with age in ME/CFS patients and controls. (e) CD8+ T cell and CD4 to CD8 T cell ratio distribution in ages older and younger than 50 years in ME/CFS and controls, (f) Correlation of NK cells with age in ME/CFS patients and controls, (g) NK cell ratio distribution in groups based on ages older and younger than 50 years. Data from healthy controls (Healthy, n = 90) and ME/CFS patients (ME/CFS, n = 186) for a (left), from Healthy (n = 91) and ME/CFS (n = 190) for a (right), b (left), c, d, and e, from Healthy (n = 91) and ME/CFS (n = 189) for b (right), f, and g, and were compared by Mann-Whitney U test for non-parametric data, with exact p values, average (AVG) and median (MED) values are shown. Correlations of data were performed using nonparametric Spearman correlation, with exact *r*_*s*_ and p value shown.

It is well established that CD4 to CD8 ratio changes are associated with aging (Yan et al., 2001;Yan et al., 2010;Serrano-Villar et al., 2014;Adriaensen et al., 2015). Indeed, CD4+ and CD8+ T cell frequencies and the CD4 to CD8 ratio correlated with age in both healthy controls (*r*_*s*_=0.4902, −0.4649, and 0.4794 respectively) and in ME/CFS subjects (*r*_*s*_= 0.4531, −0.4305, and 0.4403, respectively) (Fig. 1d). Age also showed a significant correlation with DN T cells for controls (*r*_*s*_=-0.4384), but not for ME/CFS patients (*r*_*s*_=-0.2235) (Fig. 1d). Interestingly, when we subdivided ME/CFS patients around their median age, as younger or older than 50 years of age, the difference in the percentage of CD8+ T cells and the CD4:CD8 ratio remained significant only for ME/CFS subjects younger than 50 years (p=0.0082 and 0.0131, respectively), but not for those older than 50 years (p=0.6 and 0.6, respectively) (Fig. 1e). Age also showed a significant correlation with NK cells only in ME/CFS patients (*r*_*s*_=0.3531), but not in healthy controls (*r*_*s*_=0.02794) (Fig. 1f). This change in NK cell frequency was also only seen in ME/CFS subjects younger than 50 years (p<0.0001) (Fig. 1g).

We next divided CD4+ and CD8+ T cells into naïve and memory subsets as part of their differentiation states, based on their functional and phenotypic features (Sallusto et al., 2004). To determine the proportion of these subsets in ME/CFS patients, we used well-established CD45RO and CCR7 cell surface molecules as markers for both CD4+ and CD8+ T cell subsets (Fig. 2a). Within CD4+ T cells, there was no significant difference between ME/CFS patients and healthy controls for CD45RO-CCR7+ (naïve; N) (p=0.5), CD45RO+CCR7+ (central memory; CM) (p=0.7), CD45RO+CCR7-(effector memory; EM) (p=0.2), or CD45RO-CCR7-(effector memory RA; EMRA) (p=0.06) subsets (Fig. 2b). There was also no difference in CD8+ N (p=0.4), CM (p=0.1), or EMRA (0.0509) populations, however, the CD8+ EM T cell subset was greatly increased as a proportion of CD8+ T cells (p=0.0001) in ME/CFS patients (Fig. 2c).

**Figure 2.**
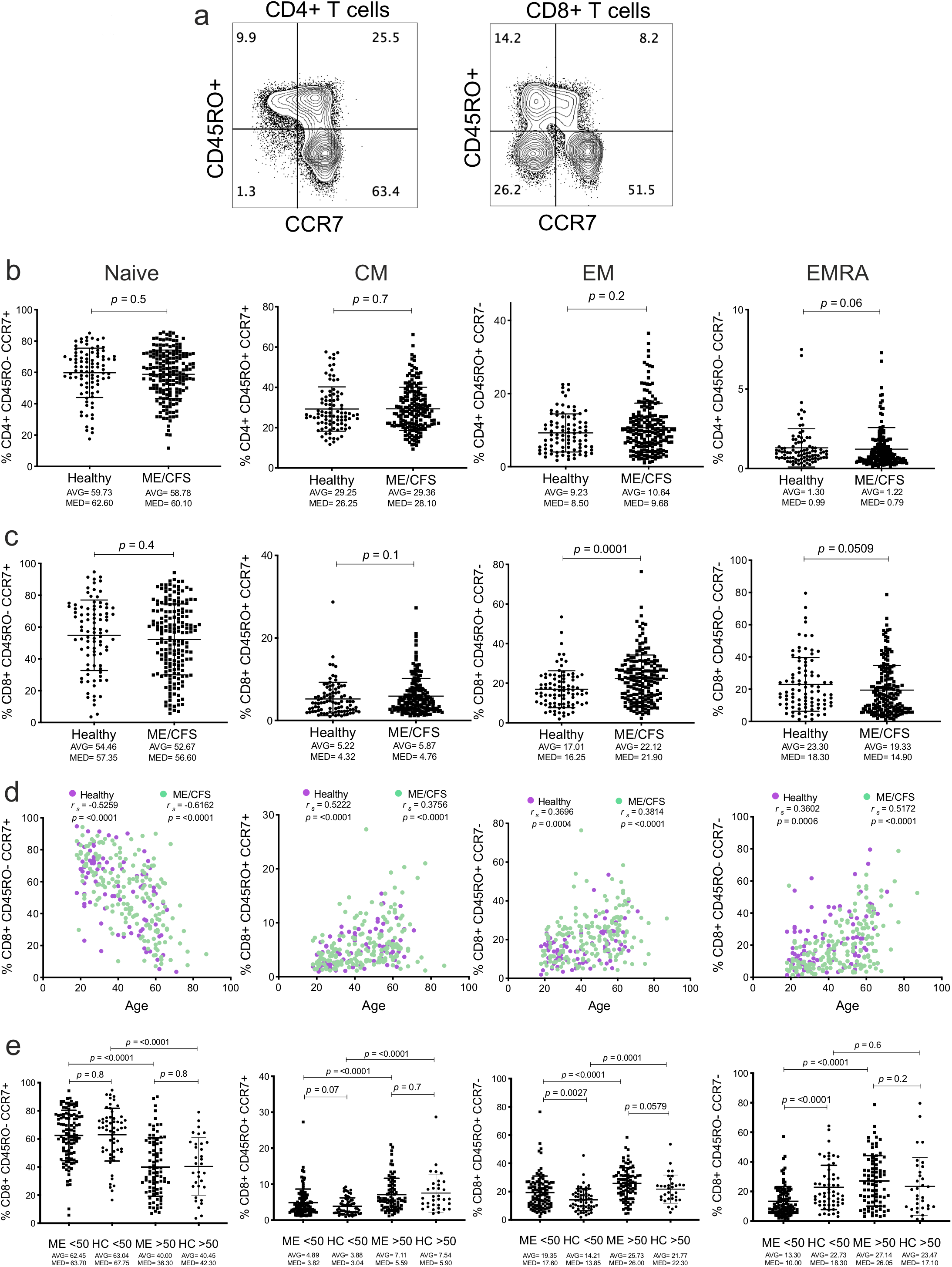
Frequencies of Naïve and memory T cell subset in ME/CFS patients. (a) T cell subsets were analyzed in PBMC with CD45RO and CCR7 expression after gating on CD3+CD4+ (left) and CD3+CD8+ T cell subsets (right), which were then subdivided into CD45RO-CCR7+ or naïve (N), CD45RO+CCR7+ or central memory (CM), CD45RO+CCR7- or effector memory (EM), and CD45RO-CCR7-effector memory RA (EMRA) subsets as shown. (b) Proportions of naïve (N), central memory (CM), effector memory (EM) and effector memory RA (EMRA) T cell subsets were analyzed within CD3+CD4+ T cells and (c) CD3+CD8+ T cells in ME/CFS and healthy subjects. (d) The frequency of each subset was correlated to subject age for CD8+ T cells, by nonparametric Spearman correlation, with exact *r*_*s*_ and p-value shown. (e) Analysis of CD8+ T cell subset frequencies in controls and ME/CFS patients that have been divided into two groups based on ages older and younger than 50 years. Data from healthy controls (Healthy, n = 91) and ME/CFS patients (ME/CFS, n = 190) for b-e, and groups were compared by Mann-Whitney test for non-parametric data, with exact p values shown, average (AVG) and median (MED) values are also shown. Correlations of data were performed using nonparametric Spearman correlation, with exact *r*_*s*_ and p value shown.

The frequencies of N, CM, EM, and EMRA populations within CD8+ T cells correlated with age for both healthy controls (*r*_*s*_=-0.5259, 0.5222, 0.3696, and 0.3602 respectively) and ME/CFS patients (*r*_*s*_=-0.6162, 0.3756, 0.3814, and 0.5172 respectively) (Fig. 2d), and this was also seen in CD4+ subsets (Supplemental Fig. 1a). However, there was no significant difference between ME/CFS patients and healthy controls for CD8+ N or CM T cells for subjects who were younger or older than 50 years (p=0.8 and 0.07, respectively) (Fig. 2d) or for CD4+ N, CM or EM subsets in different age groups of patients and controls (Supplemental Fig. 1b). Interestingly, the CD8+ EM subset difference between ME/CFS patients and healthy controls was restricted to subjects younger than 50 years (p=0.0027) (Fig. 2e). CD8+ and CD4+ EMRA subsets were also only significantly lower in ME/CFS patients who were younger than 50 years of age (p=<0.0001 and p<0.0156 respectively) (Fig. 2e and Supplemental Fig. 1b).

### Changes in Th17 cell frequency and function in ME/CFS disease

We hypothesized that ME/CFS patients may also have disruptions within effector T cell subsets resident in mucosal tissues such as Th17 cells, which respond to bacterial infections or microbiota and are also linked to autoimmune diseases (Milner et al., 2010;Pandiyan et al., 2019). To identify Th17 cells we first used CD3, CD4, CD45RO, and CCR6 expression (Fig. 3a), as almost all of this subset has a memory phenotype and also expresses the chemokine receptor CCR6 (Romagnani et al., 2009). In order to analyze the cytokine secretion from T cells, we thawed frozen aliquots of PBMC and cultured one day in IL-7 (d1) to ensure cells recovered from thawing and any dying cells could be clearly identified. We then activated the cells with PMA and Ionomycin as described in methods. The cells were then stained intracellularly for IL-17 and IFNγ and gated on cell surface expression based on CD4+CD45RO+CCR6+ and CD4+CD45RO+CCR6-cells (Fig. 3b), as previously described (Wan et al., 2011). Within the CD4+CD45RO+ (memory T cell) population, the frequency of IL-17+ (p=0.0378), IFNγ+ (p=0.0231), and IL-17+IFNγ+ (p=0.0378) secreting cells was significantly reduced in ME/CFS compared to healthy subjects (Fig. 3c).

**Figure 3.**
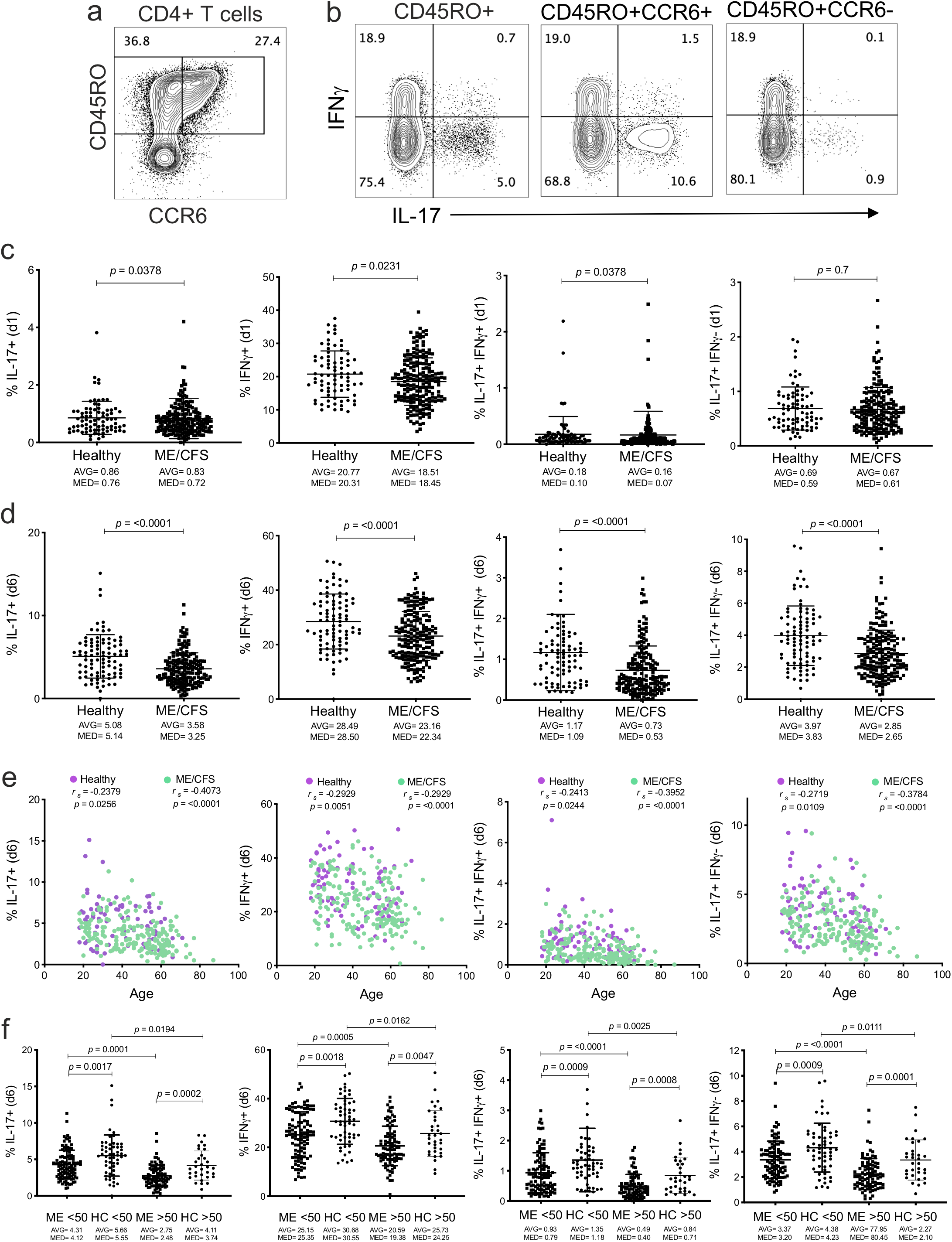
Analysis of Th17 cell frequency and function in ME/CFS subjects. PBMC purified from patient or control blood were cultured for one day (d1) in IL-7 and then stimulated and stained with specific antibodies shown for flow analysis, as described in methods. (a) CD3+CD4+ cells were gated and the proportion of CD45RO+ and CCR6+ or CCR6-cells analyzed. (b) CD4+ memory (CD45RO+) T cells expressing IFNγ and/or IL-17 or after gating into CCR6+ and CCR6-T cells. (c) The frequency of IL-17 and/or IFNγ expression in CD4+CD45RO+ memory T cells in ME/CFS patient or control PBMC. (d) Same analysis was performed in PBMC after 6-day (d6) culture in IL-7. (e) Correlation of CD4+CD45RO+ memory T cells secreting IL-17 and/or IFNγ with subject age. Groups compared by nonparametric Spearman correlation, with exact *r*_*s*_ and p-value shown. (f) Analysis of CD45RO+ memory IL-17 and/or IFNγ producing cells in control and ME/CFS patients divided into two groups based on ages older and younger than 50 years. Data from healthy controls (Healthy, n = 80) and ME/CFS patients (ME/CFS, n = 198) for c, from Healthy (n = 90) and ME/CFS (n = 195) for d, e, and f, and groups were compared by Mann-Whitney test for non-parametric data, with exact p-values, average (AVG) and median (MED) values shown. Correlations of data were performed using nonparametric Spearman correlation, with exact *r*_*s*_ and p value shown.

Previously we have shown that a portion of Th17 cells are poised to produce IL-17 or IL-22 only after priming with γc-cytokines (namely IL-2, IL-15 or IL-7) in culture, which reveal their full potential of their IL-17 secretion (Wan et al., 2011). Accordingly, we cultured PBMC from ME/CFS patients and control subjects for 6 days (d6) in IL-7 to prime Th17 cells for IL-17 secretion, as previously described (Wan et al., 2011). PBMC were then stimulated using PMA and Ionomycin, and expression of cytokines within T cell subsets was determined. In this assay, T cells from ME/CFS patients expressed profoundly lower total IL-17+ (p<0.0001), IFNγ (p<0.0001), IL-17+IFNγ+ (p<0.0001), and IL-17+IFNγ-(p<0.0001) cells (Fig. 3d), revealing a major dysfunction of Th17 cells in patients.

After 6 days in culture with IL-7, the proportion of IL-17 and IFNγ secreting cells within CD4+CD45RO+ memory population of healthy controls did not correlate with age for either IL-17+, IFNγ+, IL-17+IFNγ+, or IL-17+IFNγ-subsets (*r*_*s*_=-0.2379, −0.2929, −0.2413, and −0.2719 respectively). For ME/CFS patients, age also did not correlate with IFNγ+ expressing cells (*r*_*s*_=-0.2929), but IL-17+, IL-17+IFNγ+, and IL-17+IFNγ-subsets showed a significant correlation (*r*_*s*_=-0.4073, −0.3952, and −0.3784, respectively) (Fig. 3e). When patients were broken down into groups of subjects younger and older than 50 years, significant differences were observed in IL-17+, IFNγ+, IL-17+IFNγ+, and IL-17+IFNγ-subsets between controls and ME/CFS subjects younger than 50 years (p=0.0017, 0.0018, 0.0009, and 0.0009, respectively), as well as among the older than 50 years groups (p=0.0002, 0.0047, 0.0008, and 0.0001, respectively) (Fig. 3f).

To further investigate the disruption in the Th17 cell subset, we compared the frequency of CD4+CD45RO+CCR6+ cells between controls and ME/CFS patients. In contrast to IL-17 expression, we found that CCR6+ cells were significantly higher in ME/CFS patients (p=0.0009). However, after 6 days in IL-7, there was no difference between the subject groups (p=0.2), even though the average frequency was higher in both groups (Fig. 4a). CCR6+ cell frequency within memory CD4 T cells correlated with subject age for healthy controls (*r*_*s*_=0.3206), but not for ME/CFS patients (*r*_*s*_=0.2071). When patients were grouped as younger and older than 50 years of age, a significant difference was seen for the proportion of CCR6+ cells only in subjects younger than 50 years (p=0.0002) (Fig. 4b).

**Figure 4.**
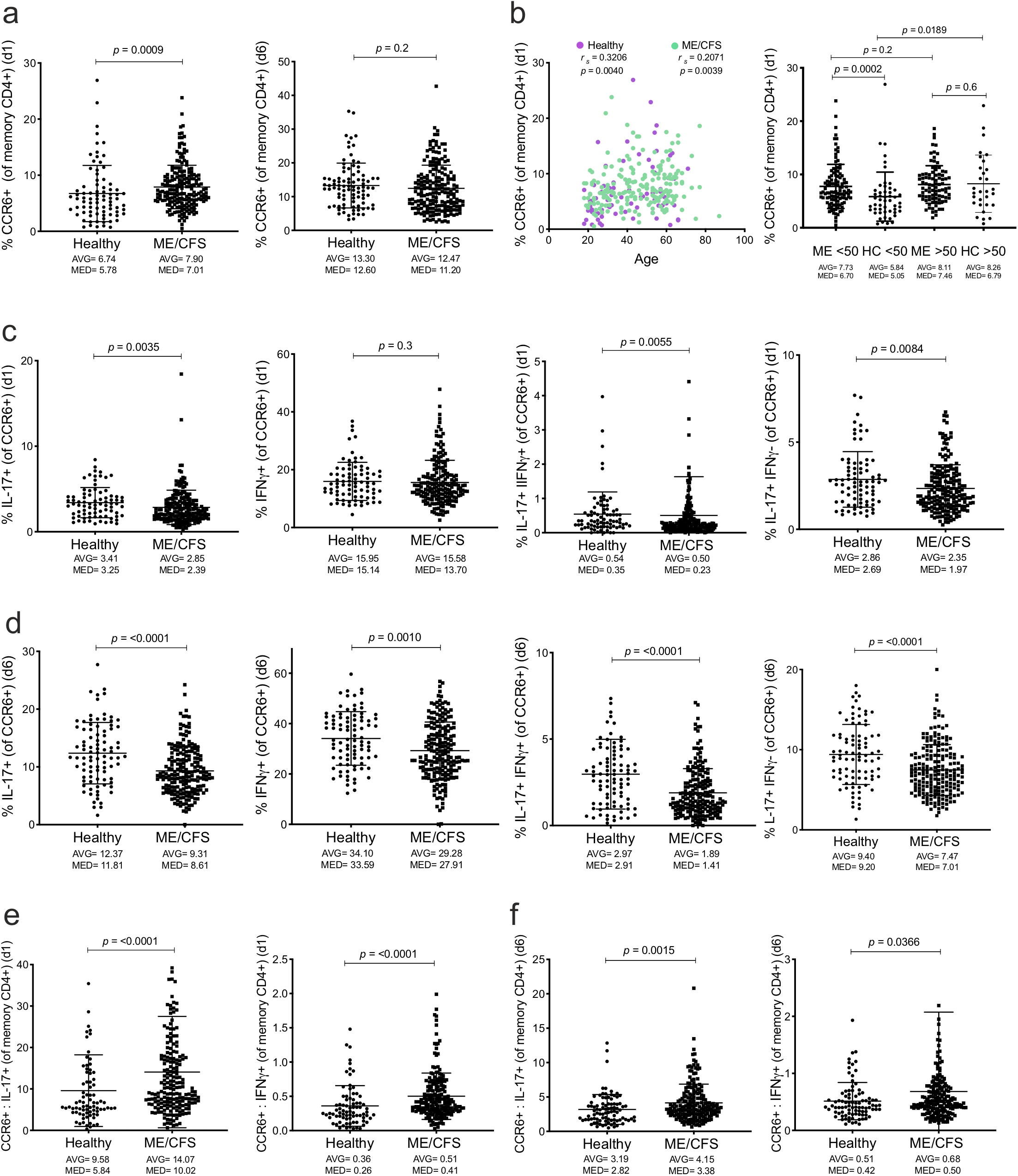
Disruption of Th17 cells in ME/CFS patients. (a) Proportion of CCR6+ T cells in memory CD4+ cells after day 1 (d1) or day 6 (d6) in culture in IL-7. (b) The frequency of CCR6+ T cells in memory CD4+ cells after day-1 culture correlated to subject age. Groups compared by nonparametric Spearman correlation, with exact *r*_*s*_ and p-value shown. Analysis of CCR6+ T cells in memory CD4+ cells after day-1 culture in healthy control and ME/CFS patients divided into two age groups, (c) IL-17 and IFNγ expression in CD4+CCR6+CD45RO+ T cells in PBMC culture in IL-7 for 1 day or (d) for 6 days, post activation as described in methods. (e) Ratio of CD4+CCR6+ cells to IL-17+ or total IFNγ+ CD4+ memory cells calculated after cells after day-1 culture or (f) after 6 days in culture with IL-7. Data from healthy controls (Healthy, n = 81) and ME/CFS patients (ME/CFS, n = 198) for a (left), from Healthy (n = 90) and ME/CFS (n = 195) for a (right), from Healthy (n = 80) and ME/CFS (n = 197) for b, from Healthy (n = 80) and ME/CFS (n = 198) for c and e, from Healthy (n = 90) and ME/CFS (n = 196) for d, from Healthy (n = 90) and ME/CFS (n = 195) for f, and groups were compared by Mann-Whitney test for non-parametric data, with exact p values shown. Average (AVG) and median (MED) are also shown. Correlations of data were performed using nonparametric Spearman correlation, with exact *r*_*s*_ and p value shown.

Remarkably, ME/CFS subjects, compared to controls, displayed lower expression of IL-17+ (p=0.0035), IL-17+IFNγ+ (p=0.0055), and IL-17+IFNγ-(p=0.0084), but not total IFNγ+ (p=0.3), within the CD4+CD45RO+CCR6+ T cells (Fig. 4c). After 6 days in culture in IL-7, the differences further increased and were seen in all cytokine-secreting cells, as a proportion of CD4+CD45RO+CCR6+ T cells, for IL-17+ (p<0.0001), IFNγ+ (p=0.0010), IL-17+IFNγ+ (p<0.0001), and IL-17+IFNγ-(p<0.0001) cells (Fig. 4d).

We next determined the ratio between the CCR6+ T cells to CD4+ memory T cells expressing IL-17 or IFNγ. Indeed, the ratio of CCR6+ cells to IL-17+ (p<0.0001) and to IFNγ+ (p<0.0001) CD4+ memory T cells were significant in ME/CFS patients compared to healthy controls (Fig. 4e). These ratios between CCR6+ cells and cytokines produced by CD4+ cells also remained higher in ME/CFS subjects after d6 in IL-7, for CCR6+ to IL-17+ cell ratio (p=0.0015), but were only marginally different for CCR6+ to IFNγ+ cell ratio (p=0.0366) (Fig. 4f).

We have previously shown that CD161 within the CD4+CD45RO+CCR6+ T cells can further divide these cells into subsets with differences in IL-17 and IFNγ secretion (Wan et al., 2011). As such, we further divided CCR6+ cells based on CD161 expression (Fig. 5a). The proportion of CD161+ cells within the CCR6+ subset was only slightly different in ME/CFS compared to controls (p=0.0439) (Fig. 5a). We then analyzed IL-17 and IFNγ expression within the CD161+ and CD161-subsets of CD4+ CD45RO+ CCR6+ cells after 6 days in culture (Fig. 5b). Within the CD4+CCR6+CD161+ cells, there was a significant difference in expression of IL-17+, IL-17+IFNγ+, and IL-17+IFNγ-cells between ME/CFS and controls (p<0.0001 for all), but not for IL-17-IFNγ+ cells (p=0.06) (Fig. 5c). CD161-cells within the CD4+CD45RO+CCR6+ subset also displayed lower IL-17+, IL-17+IFNγ+, IL-17+IFNγ-, and IL-17-IFNγ+ in ME/CFS subjects compared to healthy controls (p=<0.0001, <0.0001, 0.0001, and 0.0042, respectively), (Fig. 5d).

**Figure 5.**
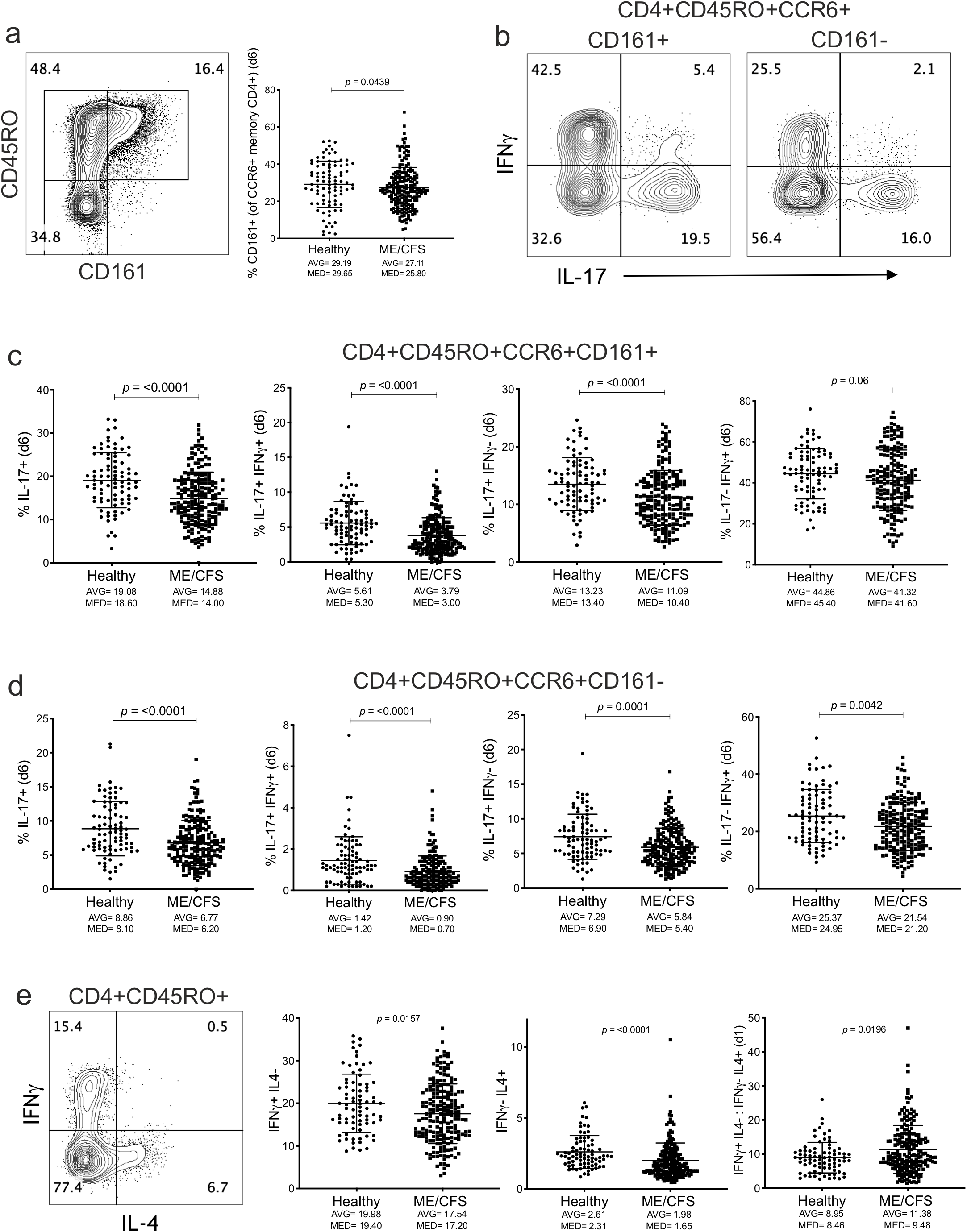
Disruption of Th17 cells. PBMC from ME/CFS patients and healthy controls were cultured in IL-7 for 6 days and stimulated with PMA/ionomycin for 4 hours as described in methods. (a) Live CD4+ T cells were gated on different memory subsets based on CD161 expression (left) and the proportion of CD161+ cells within CD4+CD45RO+CCR6+ subset is shown for each subject (right). (b) Intracellular expression of IL-17 and IFNγ within CD4+CCR6+CD161+ and CD4+CCR6+CD161-T cell subsets. (c) The frequencies of IL-17 and IFNγ expressing cells within CD4+CD45RO+CCR6+CD161+ and (d) CD4+CD45RO+CCR6+CD161-cells were calculated and shown for individual study participants. (e) Analysis of IFNy+IL4-(Th1 cells) and IFNγ-IL4+ (Th2 cells) in memory CD4+ cells after day 1 (d1) in culture in IL-7, and ratio of Th1 to Th2 cells. Data from healthy controls (n = 90) and ME/CFS patients (n = 196) for a (right), from Healthy (n = 87) and ME/CFS (n = 191) for c and d, from Healthy (n = 90) and ME/CFS (n = 198) for e, and groups were compared by Mann-Whitney test for non-parametric data, with exact p-values and average (AVG) and median (MED) values are shown. Correlations of data were performed using nonparametric Spearman correlation, with exact *r*_*s*_ and p value shown.

In CD4+ memory T cells, in addition to IL-17 expression, we also determined the frequency of T cells that were either expressing IFNγ (IFNγ+IL-4-) or IL-4 (IFNγ-IL-4+) only, which respectively define Th1 and Th2 T cell subsets (Fig. 5e). We found that the proportion of IFNγ+IL-4- and IFNγ-IL-4+ within CD4+ memory T cells was significantly lower (p=0.0157 and p<0.0001 respectively) in ME/CFS subjects (Fig. 5e). However, the ratio of Th1 (IFNγ+IL-4-) to Th2 (IFNγ-IL-4+) was higher in ME/CFS patients compared to the control group (p=0.0196) (Fig. 5b), suggesting an imbalance of Th1 to Th2 cells. Together, these findings highlight major functional perturbations within the CD4+ T cell subset in the ME/CFS patient cohort.

### Changes in Frequency of MAIT cells in ME/CFS

Mucosal-associated invariant T (MAIT) cells are a subset of the non-classical T cell population and defined by an invariant T cell receptor that is triggered by riboflavin metabolites produced by bacteria, including commensal microbiota (Tastan et al., 2018;Godfrey et al., 2019). Similar to the Th17 subset, we hypothesized that dysbiosis in the gut microbiome or prior bacterial infections may result in changes in MAIT cell frequencies or function. To identify MAIT cells in PBMC, we used Vα7.2 and CD161 surface molecules as previously described (Khaitan et al., 2016;Tastan et al., 2018). We then determined the frequency of MAIT cells within CD4+, CD8+ and CD4-CD8-(double negative or DN) T cell compartments in ME/CFS patients and healthy controls (Fig. 6a). There was no significant difference between patients and controls for CD4+ (p=0.7), CD8+ (p=0.7), or double negative (DN) MAIT cells (p=0.2) as a proportion of the CD4+, CD8+ and DN T cells respectively (Fig. 6b). However, CD4+ and CD8+ MAIT cell frequencies in PBMC after 6-day culture in IL-7 showed a significant difference (p=0.0250 and p=0.0221 respectively) between ME/CFS patients and controls, but DN MAIT cell frequency did not change between ME/CFS and control samples after 6 days culture (p=0.3) (Fig. 6c). When the ratio of MAIT cell frequency at day 0 (d0) vs day 6 after IL-7 culture (d6) was assessed, we found that the frequency of CD8+ MAIT cells in ME/CFS PBMC was greatly reduced after 6 days of culture compared to d0 levels (p=0.0008), but there was no significant difference seen for CD4+ (p=0.06) or for DN MAIT cells (p=0.8) between ME/CFS patients and controls (Fig. 6d). Corollary to this finding, the ratio of CD8+ MAIT to DN MAIT cells in ME/CFS patients and controls was only slightly significant at d0 (p=0.0364), but became highly significant after 6 days in culture with IL-7 (p<0.0001) (Fig. 6e). Together, these findings suggest that CD8+ MAIT cells from ME/CFS subjects survived less in in vitro culture with IL-7.

**Figure 6.**
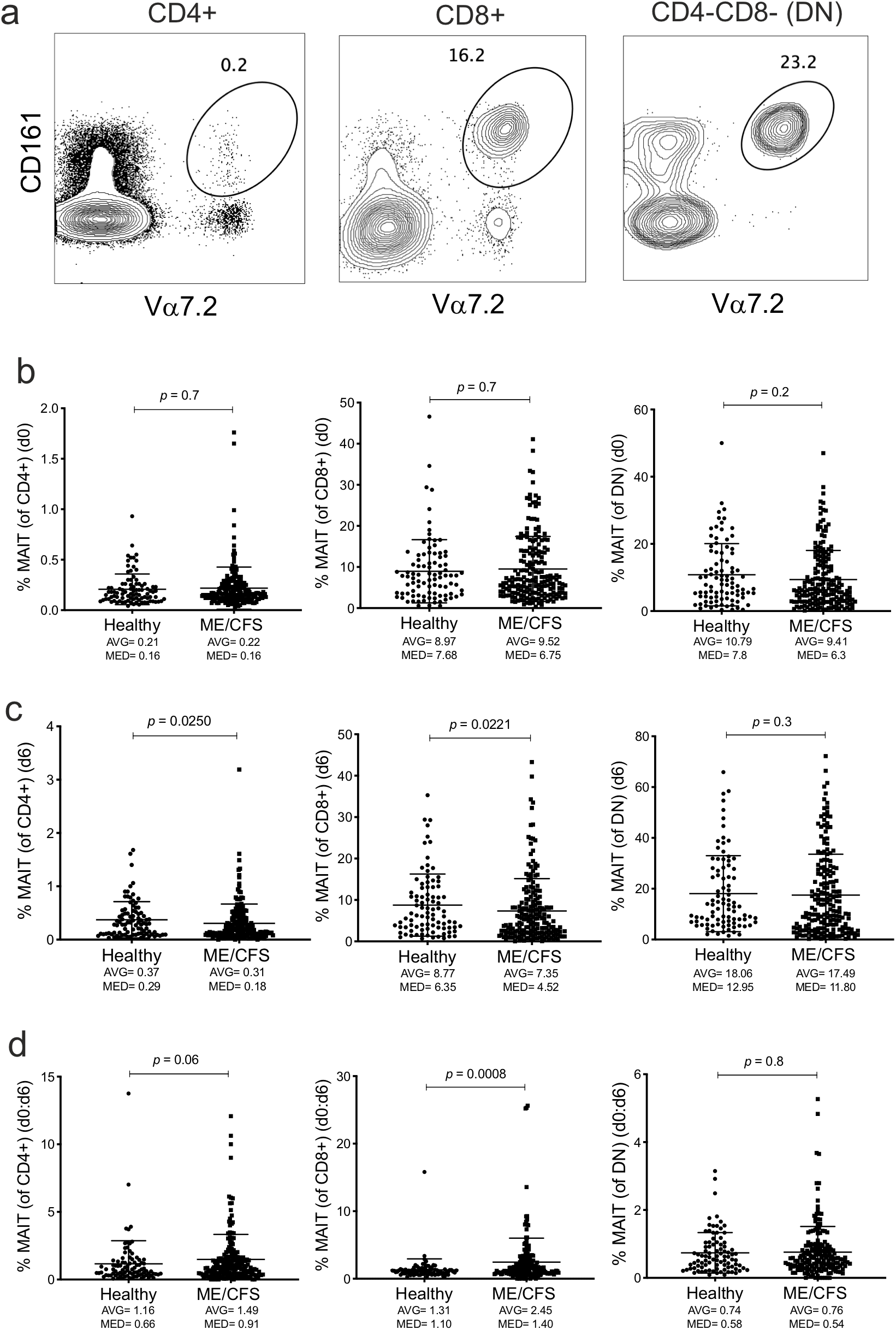
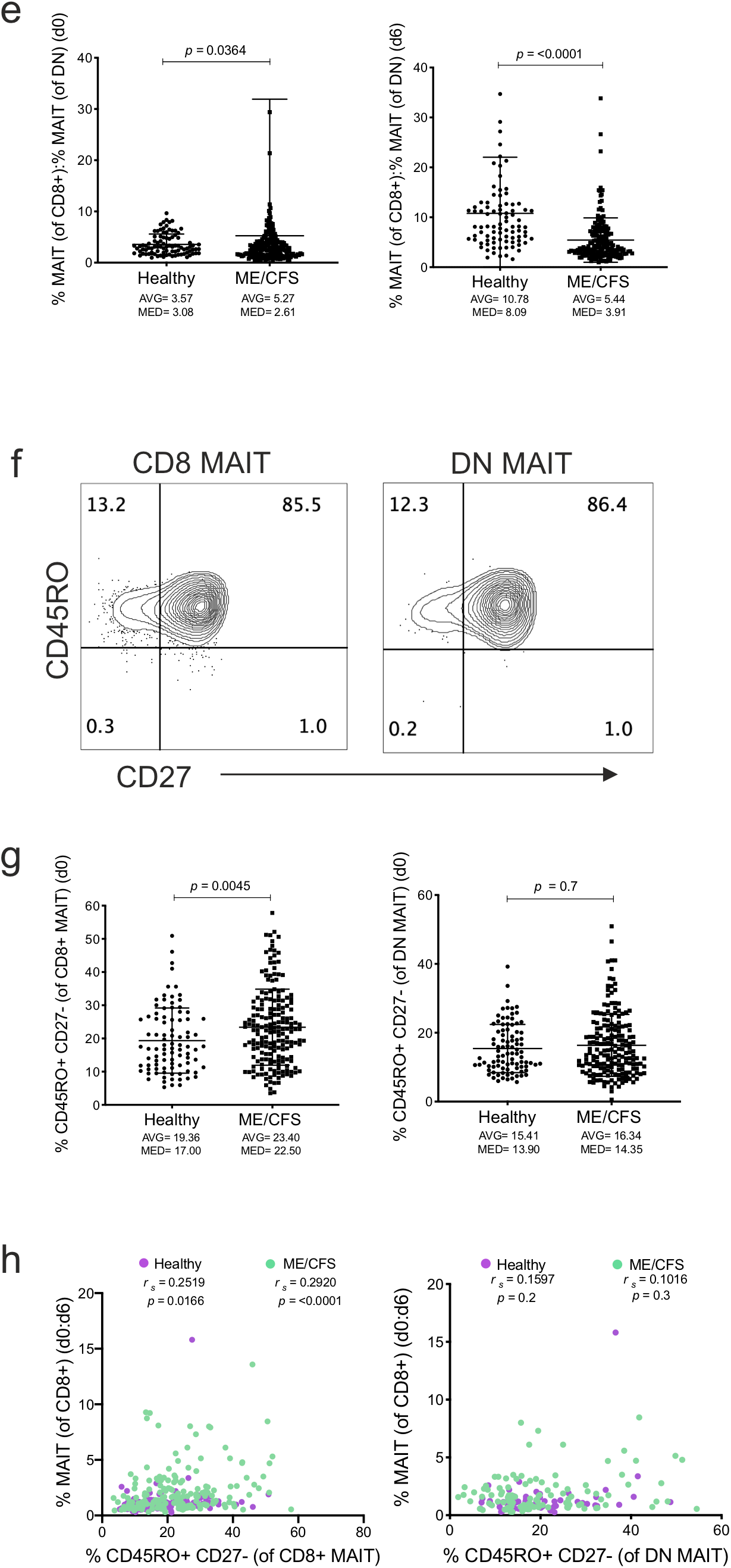
Perturbation of Mucosal Associated Invariant T (MAIT) cell subsets in ME/CFS PBMC. (a) MAIT cell subset frequencies were identified based on the co-expression of CD161 and Vα7.2, after gating within CD4+, CD8+ and CD4-CD8-(DN) T cells (b) Analysis of the proportion of MAIT cells in each of these T cell subsets on day 0 (d0) and (c) 6 days (d6) in culture with IL-7. (d) Ratio of day 0 to day 6 MAIT cells was calculated for individual study participants. (e) The ratio of CD8+ MAIT to DN MAIT cells was calculated for day 0 (left) and day 6 (right). (f) Surface expression of CD27 was determined after gating for CD8+ MAIT and DN MAIT cell subsets as shown. (g) Analysis of the proportion of CD45RO+CD27-within CD8+ MAIT and DN MAIT cells in PBMC of ME/CFS and control subjects. Groups were compared by Mann-Whitney test for non-parametric data, with exact p values, average (AVG) and median (MED) values are shown. (h) The ratio of MAIT cell subset (CD8+ or DN separately) frequency at day 0 to day 6, was correlated with CD27-MAIT cell frequency (of total CD8+ MAIT cells). Data from healthy controls (Healthy, n = 91) and ME/CFS patients (ME/CFS, n = 190) for b, from Healthy (n = 90) and ME/CFS (n = 195) for c (left and middle), from Healthy (n = 90) and ME/CFS (n = 196) for c (right), from Healthy (n = 90) and ME/CFS (n = 186) for d (left), from Healthy (n = 90) and ME/CFS (n = 190) for d (middle), from Healthy (n = 90) and ME/CFS (n = 189) for d (right), from Healthy (n = 91) and ME/CFS (n = 190) for e (left), from Healthy (n = 90) and ME/CFS (n = 196) for e (right), from Healthy (n = 91) and ME/CFS (n = 190) for g, from Healthy (n = 90) and ME/CFS (n = 184) for h (left), from Healthy (n = 60) and ME/CFS (n = 108) for h (right), and groups were compared by Mann-Whitney test for non-parametric data, with exact p-values and average (AVG) and median (MED) values are shown. Correlations of data were performed using nonparametric Spearman correlation, with exact *r*_*s*_ and p value shown.

Because CD27 expression on MAIT cells could indicate a recently activated or differentiated subset, similar to other CD8 T cells (Dolfi and Katsikis, 2007;Grant et al., 2017), we evaluated CD27 expression in MAIT subsets (Fig. 6f). We found that ME/CFS patients had a significant difference where there were higher CD45RO+CD27-cells compared to the control group (p=0.0045), but interestingly, this difference was not observed within DN MAIT cells (p=0.7) (Fig. 6g). The d0 to d6 CD8+ MAIT cell ratio also displayed a slight positive correlation with the frequency of CD27-CD8+ MAIT cells in patients (*r*_*s*_=0.2920), but not in healthy controls (*r*_*s*_=0.2519). In contrast, CD27-DN MAIT cells did not correlate with the d0 to d6 cell frequency ratio for either controls (*r*_*s*_=0.1597), or ME/CFS patients (*r*_*s*_=0.1016) (Fig. 6h).

We then asked to what extent MAIT cells were functionally different between ME/CFS patients and controls. For this approach we first stimulated the PBMC with a cocktail of three cytokines, namely IL-12+IL-15+IL18, since this combination has been uniquely shown to induce expression of IFNγ from MAIT cells (Ussher et al., 2014;Salou et al., 2017). Accordingly, IFNγ along with Granzyme A expression was used to evaluate the response of CD8+ MAIT and CD8+ non-MAIT cells in PBMC to stimulation with IL-12+IL-15+IL18 cocktail (Fig. 7a). ME/CFS patient PBMC stimulated with the cytokine cocktail showed much lower IFNγ+ MAIT cells (p=<0.0001), but induction of IFNγ+ from non-MAIT CD8+ T cells was comparable (p=0.1) to healthy subjects (Fig. 7b). Granzyme A expressing MAIT cells were also much higher in ME/CFS subjects (p<0.0001), but non-MAIT cells expressing Granzyme A were not different (p=0.8) between controls and patients (Fig. 7c). In addition, CD27-CD8+ MAIT cells and IFNγ+ MAIT cells upon cytokine stimulation were negatively correlated in ME/CFS patients (*r*_*s*_=-0.3431) but not in controls (*r*_*s*_=-0.2112) (Fig. 7d). CD27-CD8+ MAIT cells were not correlated with CD8+ non-MAIT IFNγ+ cells for either healthy controls (*r*_*s*_=-0.1370) or ME/CFS patients (*r*_*s*_=-0.09816) (Fig. 7d).

**Figure 7.**
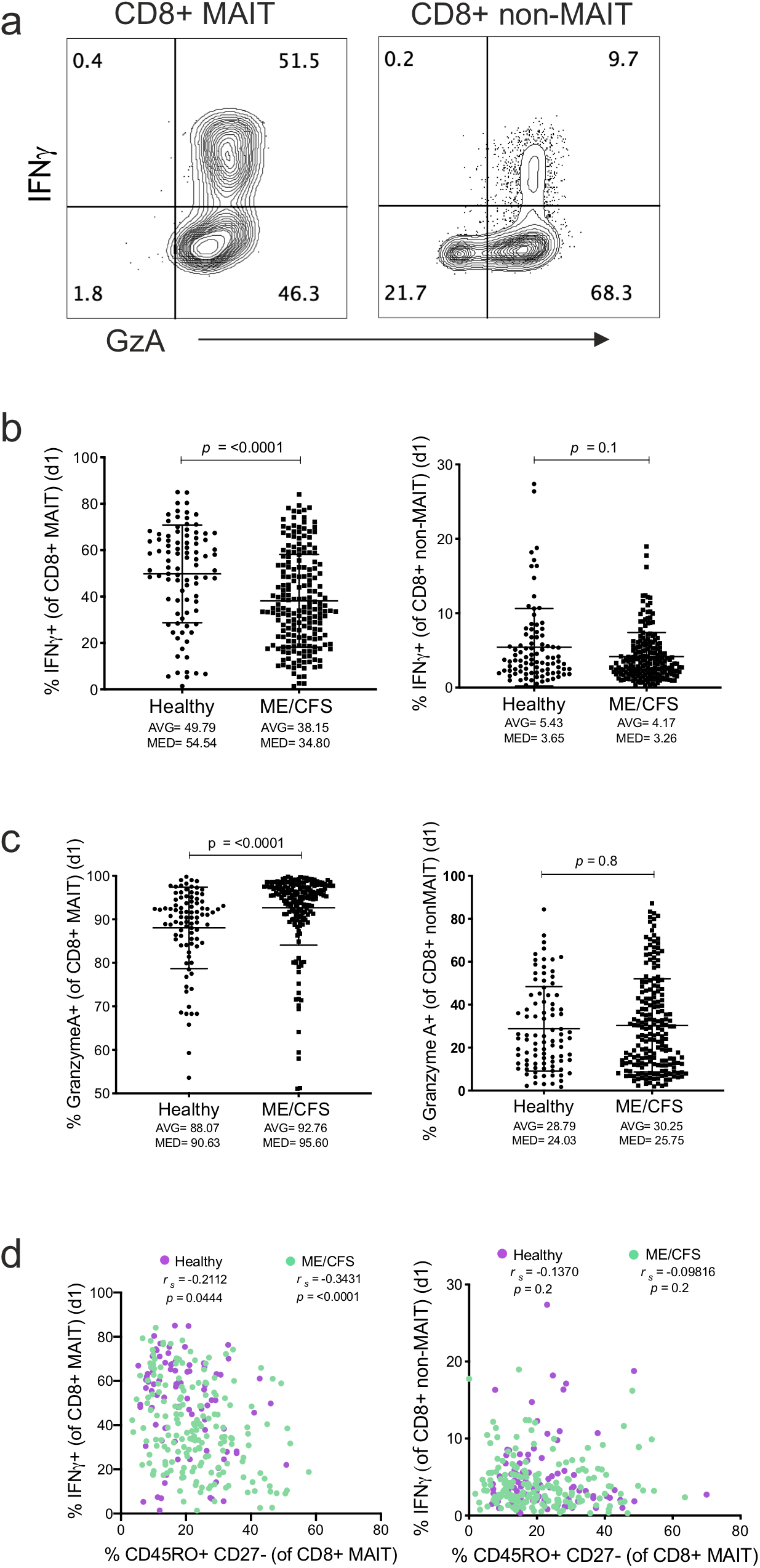
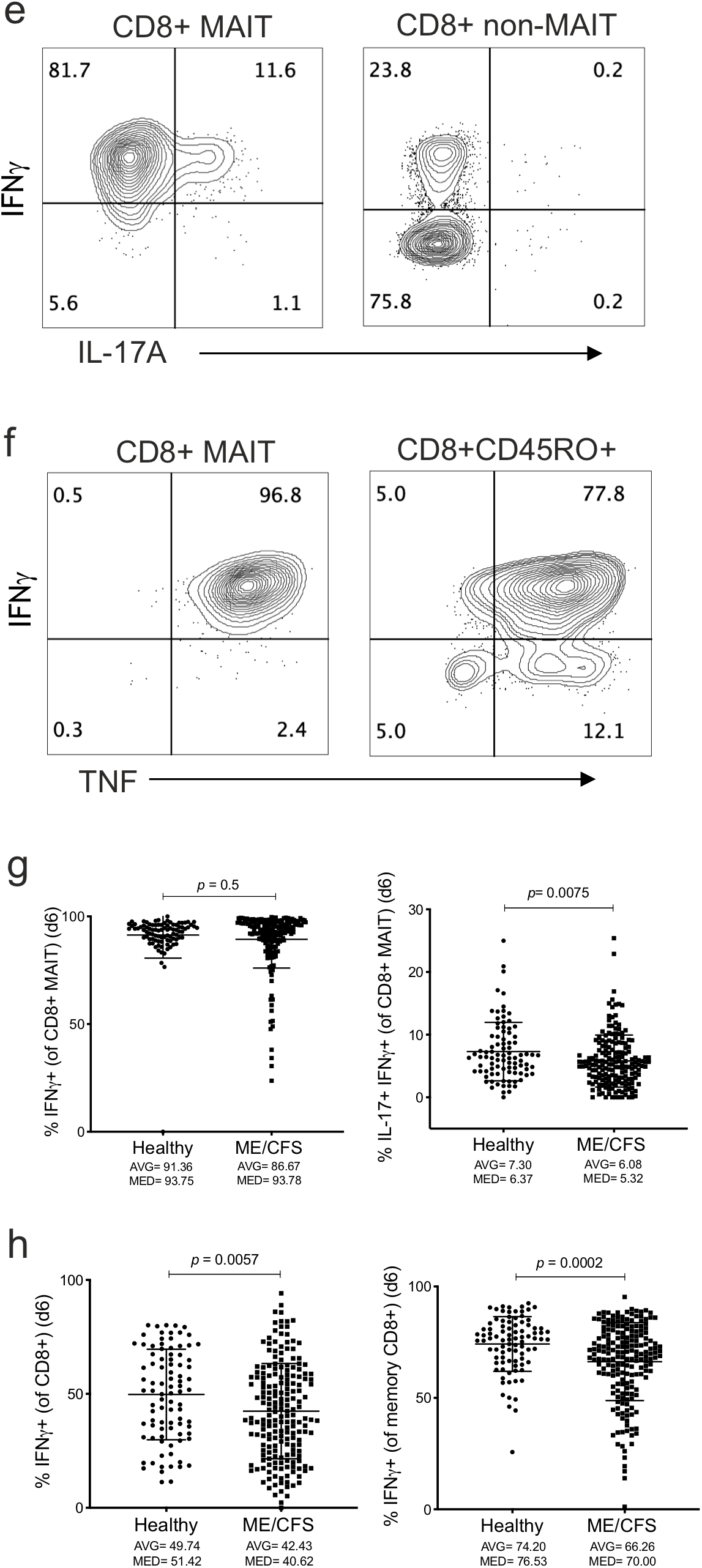
Activation of MAIT cell effector functions. (a) PBMC were stimulated with combination of the cytokines IL-12+IL-15+IL-18 for 1 day as described in methods, and intracellularly stained for IFNγ and Granzyme A expression, which was analyzed after gating on MAIT (CD161+Vα7.2+) and non-MAIT (CD161-Vα7.2-) CD8+ T cells. (b) Proportion of IFNγ and (c) Granzyme A in CD8+ MAIT and non-MAIT cells from ME/CFS and control subjects. (d) The frequency of CD8+CD45RO+CD27-MAIT cells was correlated to CD8+ MAIT and non-MAIT IFNγ+ cells after stimulation with cytokine combination. Groups were compared by nonparametric Spearman correlation, with exact *r*_*s*_ and p value shown in figures. (e) PBMC were cultured in IL-7 for 6 days (d6) then stimulated with PMA and Ionomycin as described in methods. Frequency of IFNγ and IL-17A expression within MAIT and non-MAIT CD8+ T cells were compared between patient and control groups. (f) IFNγ and TNFα expression, after activation, in CD8+ MAIT, and CD8+CD45RO+ non-MAIT memory T cells. (g) Expression of IFNγ or IL-17+IFNγ cells within CD8+ MAIT cells in ME/CFS and control subjects. (h) Proportion of CD8+ or CD8+ memory (gated on CD45RO+) cells expressing IFNγ. Data from healthy controls (Healthy, n = 91) and ME/CFS patients (ME/CFS, n = 198) for b and c, from Healthy (n = 91) and ME/CFS (n = 185) for d (left), from Healthy (n = 91) and ME/CFS (n = 191) for d (right), from Healthy (n = 91) and ME/CFS (n = 188) for g (left), from Healthy (n = 90) and ME/CFS (n = 183) for g (right), from Healthy (n = 90) and ME/CFS (n = 196) for h, and groups were compared by Mann-Whitney test for non-parametric data, with exact p-values, average (AVG) and median (MED) values are shown. Correlations of data were performed using nonparametric Spearman correlation, with exact *r*_*s*_ and p value shown.

Since MAIT cells have also been shown to express IL-17, similar to Th17 cells (Salou et al., 2017), we next sought to determine the production of IL-17 and IFNγ from MAIT cells in response to PMA and Ionomycin stimulation. There was very little to undetectable IL-17 expression from MAIT cells after one day in culture (data not shown). However, after 6 days in IL-7, MAIT cells expressing IL-17 were greatly increased upon PMA and Ionomycin stimulation, however, IL-17 remained undetectable in non-MAIT CD8+ T cells (Fig. 7e). This finding suggests that MAIT cells can also undergo priming with cytokines, similar to the classic Th17 cells (Wan et al., 2011) and IL-17 expression mimics tissue-resident MAIT cells (Sobkowiak et al., 2019).

In addition, we also determined IFNγ and TNFα secretion from CD8+ MAIT and CD8+ non-MAIT CD45RO+ (memory) T cells after 6 days in culture with IL-7 (Fig. 7f). There was no significant difference in ME/CFS patients for IFNγ+ MAIT cells (p=0.5), but we found highly reduced IL-17+IFNγ+ MAIT cells in ME/CFS patients compared to healthy controls (p=0.0075) (Fig. 7g). The frequency of IFNγ+ secreting cells was also reduced within CD8+ non-MAIT cells (p=0.0057) and within CD8+CD45RO+ memory T cells (p=0.0002), in ME/CFS PBMC cultured for 6 days in IL-7 (Fig. 7h).

### Changes in regulatory T (Treg) cells in ME/CFS patients

Regulatory T (Tregs) cells are critical in controlling autoreactive or excessive immune responses. Given the observed perturbance in the effector functions of T cell subsets that suggest chronic immune activation, we hypothesized that there would be a corresponding increase in Tregs in ME/CFS patients. For this experiment, we used Foxp3 and Helios as markers to assess Treg cell frequencies within both naïve and memory CD4+ T cells, as previously described (Mercer et al., 2014) (Fig. 8a). Indeed, frequencies of both naïve Tregs (p=0.0005), and memory Tregs (p=0.0094) were increased in ME/CFS compared to controls (Fig. 8b).

**Figure 8.**
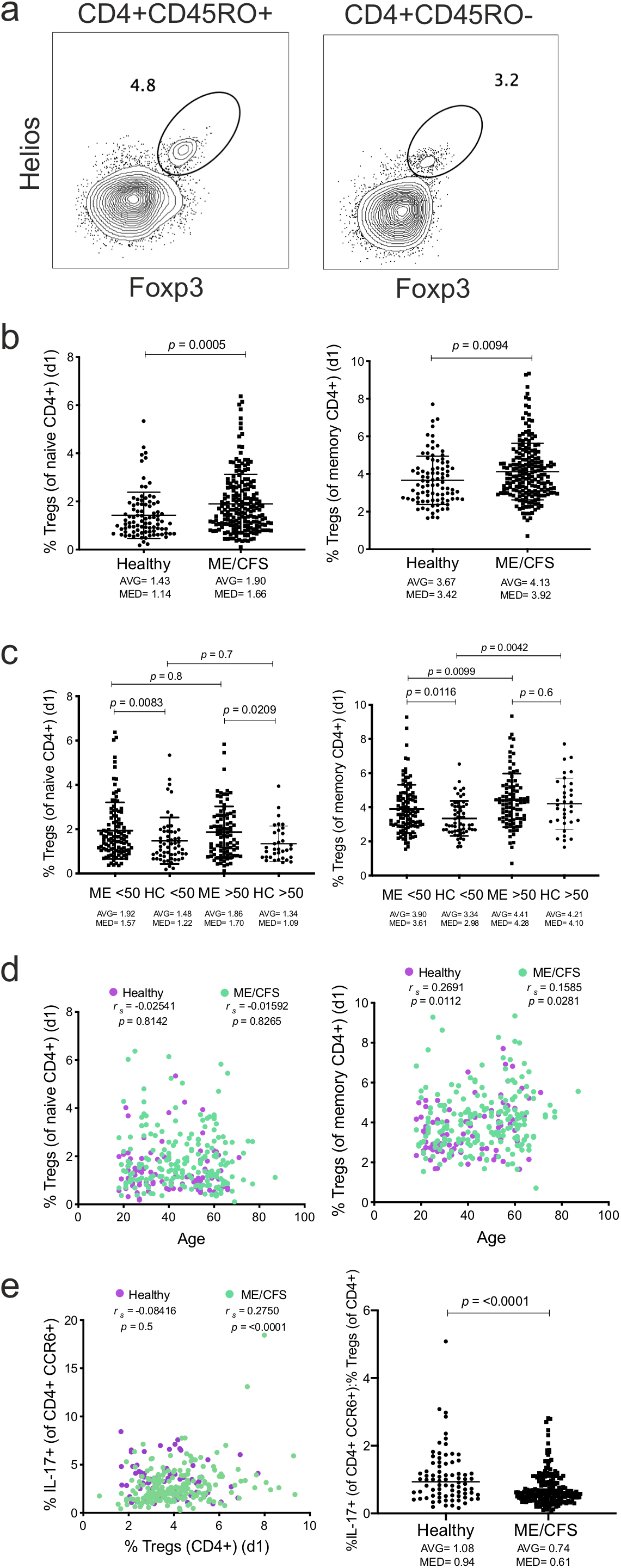
Regulatory T (Treg) cell frequency in ME/CFS. (a) PBMC were stained with Foxp3 and Helios intracellularly and expression was analyzed after gating on CD4+ naïve (CD27+CD45RO-) and memory (CD45RO+) T cells as described in the methods. (b) Proportions of Tregs (Foxp3+Helios+) were calculated within CD4+ naïve and memory subsets in ME/CFS and healthy subjects. (c) CD4+ naïve and memory Tregs divided into two groups based on age younger and older than 50 years in all subjects. (d) Correlation of CD4+ naïve and memory Treg subset frequencies with subject age. (e) Correlation between Th17 cells and memory Tregs was performed by nonparametric Spearman correlation, with exact *r*_*s*_ and p-value shown. The ratio of IL-17-expressing cells within Th17 subset (CCR6+) to memory Tregs were compared between ME/CFS subjects and controls. Data from healthy controls (Healthy, n = 91) and ME/CFS patients (ME/CFS, n = 197) for b, c, and d, from Healthy (n = 80) and ME/CFS (n = 197) for e, and groups compared by the Mann-Whitney test for non-parametric data, with exact p value, average (AVG) and median (MED) values are shown. Correlations of data were performed using nonparametric Spearman correlation, with exact *r*_*s*_ and p value shown.

When broken down into groups where subjects were younger or older than 50 years, naïve Tregs showed a highly significant difference in ME/CFS patients vs controls in the younger than 50 years group (p=0.0083), and a slightly significant difference in ME/CFS patients vs controls in the older than 50 years group (p=0.0209). The difference in memory Tregs was also significant between ME/CFS patients and controls younger than 50 years (p=0.0116), but not when older than 50 years groups were compared (p=0.6) (Fig. 8c). There was no correlation with subject age for ME/CFS patients or controls for naïve Tregs (*r*_*s*_=-0.02541 and *r*_*s*_=-0.01592, respectively), or for memory Tregs (*r*_*s*_=0.2691 and *r*_*s*_=0.1585, respectively) (Fig. 8d).

The ratio of Th17 cells to Tregs is an important feature that is perturbed during chronic inflammatory conditions or autoimmune diseases. Therefore, we also determined this ratio in ME/CFS patients vs healthy controls. While the Th17 (CCR6+ IL-17-secreting cells) frequency did not correlate with memory Treg cells in ME/CFS patients (*r*_*s*_=0.2750) or healthy controls (*r*_*s*_=-0.08416), remarkably, the ratio of these two related subsets were also highly different between the ME/CFS patients and the healthy controls (p<0.0001) (Fig. 8e)

### Machine learning analysis to identify predictive immune parameters for ME/CFS

Our immune profiling analysis identified many T cell subset parameters that were different in ME/CFS patients vs healthy controls. From the total of 65 immune profile features, 40 features were identified as different at a 5% false discovery rate (Supplemental Table 2). However, while some of these were highly significant, given the high variability and ranges in humans for immune parameters, on their own they would not have clinically relevant specificity and sensitivity to discriminate patients from healthy individuals. Therefore, we decided to use a classifier model using a machine learning algorithm called the random forest (RF) classifier (Wang and Li, 2017).

The RF classifier or algorithm is an ensemble method that depends on a large number of individual classification trees (Wang and Li, 2017;Huynh-Thu and Geurts, 2019). Each classification tree emits a predicted class and the class with the most votes becomes the model prediction. The individual trees are designed using a randomly selected number of samples (sampling with replacement) and a randomly selected feature set to minimize correlation between trees. A large number of relatively uncorrelated classification trees (models) are combined to provide a robust classification of the individual sample (Aevermann et al., 2018). As such, we implemented an RF model to classify ME/CFS patients and healthy controls using the immune profiling data. The performance of the RF was evaluated using the receiver operating characteristic (ROC) curve, which is created by plotting the true positive rate (TPR) against the false positive rate (FPR). The class prediction probability of a sample can be computed based on the proportion of votes obtained for that call. Given a threshold T for the probability, a sample is classified as an ME/CFS patient if the probability is higher than T and the ROC curve plots TPR against the FPR.

The area under the ROC curve which is denoted by AUC is equal to the probability that a randomly chosen positive instance will be ranked higher than a randomly chosen negative instance. A perfect classifier will have the maximal area under the curve of 1. The ROC curves of the RF classifier corresponding to 4 subsets of immune profile features are shown in Figure 9. The AUC of the RF classifier using all 65 features is ~0.93, meaning that there is a chance of 93% that the classifier will correctly distinguish between patients and healthy controls (Fig. 9 and Table 1). The 40 significantly different features or top 10 features with the highest importance score among these 40 significantly different immune parameters had slightly lower AUC scores (~0.92 and ~0.88, respectively), whereas the top 10 significantly different features had a lower AUC score (~0.82) (Fig. 9 and Table 1, full list of the features are in supplemental Tables 2, 3).

**Figure 9.**
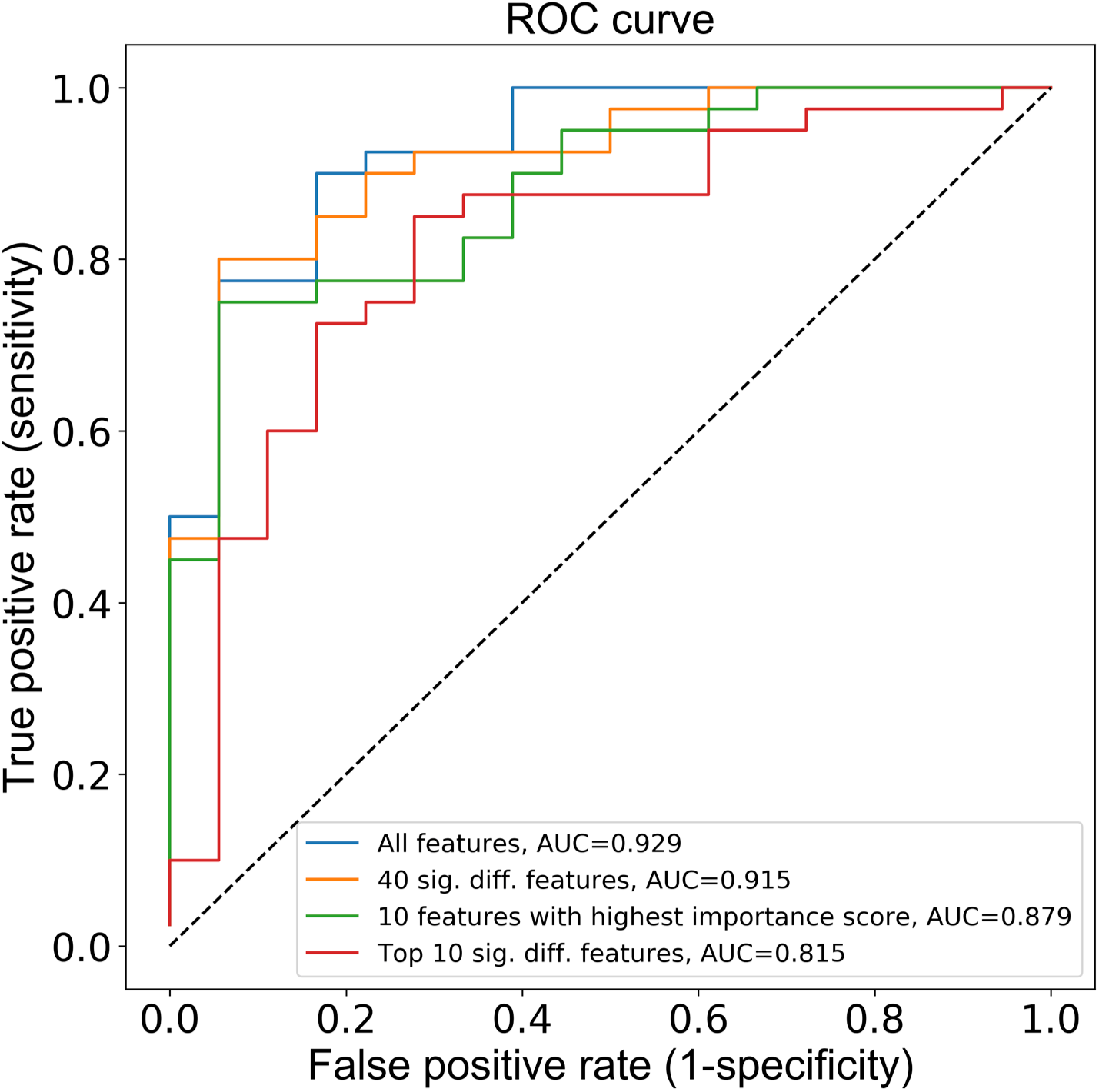
Random forest clustering of immune features in ME/CFS and control subjects. To generate a receiver operating characteristic (ROC) curve using random forest (RF) clustering algorithm, a training set with 231 samples (80% of total samples) was selected and the remaining data, corresponding to 58 samples (20% of total samples), was left as the test set. Missing values in the training and test sets were replaced by the corresponding median value in the training set. A K-fold cross-validation method was used (K=3) to tube the hyperparameters of the model and was trained using a distinct set of features as input; all 65 immune profile features, the 40 significantly different features, the top 10 significantly different features and the top 10 features that received the highest importance score are plotted.

**Table I:** Metrics of the RF classifier using different numbers of immune profile features. The sensitivity (recall), specificity, positive predictive value (precision), negative predictive value, accuracy and F_l_ score of immune features are shown. Detailed explanation of these metrics and formulas to calculate them are given in methods section. The rows represent the metrics of: 1) RF classifier when all 65 features are used, 2) for 40 significantly different immune features, 3) the 10 features with the highest importance score among the 40 significantly different immune parameters, and 4) the RF classifier when the top 10 significantly different features are used.

| Number of features | Sensitivity | Specificity | Positive pred. value | Negative pred. value | Accuracy | $F_1$ score | AUC |
| --- | --- | --- | --- | --- | --- | --- | --- |
| 65 total set | 0.950 | 0.611 | 0.844 | 0.846 | 0.845 | 0.894 | 0.929 |
| 40 significant | 0.900 | 0.722 | 0.878 | 0.765 | 0.845 | 0.889 | 0.915 |
| 10 important | 0.925 | 0.556 | 0.822 | 0.769 | 0.810 | 0.871 | 0.879 |
| 10 significant | 0.825 | 0.722 | 0.868 | 0.650 | 0.793 | 0.846 | 0.815 |

## Discussion

Several studies of the immune system of ME/CFS subjects have revealed disruptions in the number and function of T cell subsets, B cells and natural killer (NK) cells in patient blood (Fletcher et al., 2010;Brenu et al., 2012;Curriu et al., 2013;Brenu et al., 2014). Here we extend these findings and show a highly significant disruption in several key T cell subset frequencies, and importantly, their effector functions. Our findings reveal profound changes in the functional capacity of mucosal associated T cell (MAIT) and Th17 cells, Tregs, and signs of CD8+ T cell and NK cell disruption in a subset of younger subjects. Given that these cell types are involved in regulating viral or bacterial infections or the microbiota, our findings suggest that some of the ME/CFS patients have severe immune perturbations likely triggered by chronic infections or are associated with changes in the microbiome.

Our observation that both CD8+ T cells and NK cells are proportionately reduced in ME/CFS subjects may also reflect chronic viral infection or persistence (such as CMV or EBV infections) that triggers a chronic activated state of these cells. It is important to note that this difference was predominantly observed in subjects younger than 50 years old, as healthy subjects over the age of 50 also begin to display a similar phenotype as patients and thus were no longer statistically different compared to patient subjects. Indeed, this is consistent with the increase in chronic viral infections seen during normal aging, such as CMV seropositivity (Ellefsen et al., 2002). However, given this distribution, these subset differences alone are unlikely to be the main cause of ME/CFS symptoms as it is apparent there are also multiple and severe changes in the effector functions of other T cell subsets. In addition, the proportion of CMV/EBV infection among healthy controls and ME/CFS patients was not significantly different in either age group (data not shown).

Another T cell subset that we found to be highly disrupted in ME/CFS subjects was Th17 cells, which are characterized by secretion of IL-17, expression of chemokine receptor CCR6 and transcription factor RORC (Sallusto et al., 2012). Th17 cells play a major role in both immune response to bacterial or fungal infections and in pathogenesis of autoimmune or chronic inflammatory diseases (Tesmer et al., 2008;Sallusto and Lanzavecchia, 2009). We previously found that a significant portion of cells programmed towards the Th17 phenotype do not immediately secrete IL-17 but require a priming stage with IL-7 or IL-15, which we call poised Th17 cells (Wan et al., 2011). Indeed, when we performed an assay by culturing PBMC in IL-7 for 6 days before analyzing the IL-17 production from these poised CCR6+ Th17 cells, the differences between ME/CFS patients and controls became profound. However, while there were less Th17 cells producing IL-17 in ME/CFS patients, paradoxically the proportion of CCR6+ Th17 cells were significantly higher compared to controls. Together these findings suggest chronic activation of the Th17 subset that potentially induces an “exhausted” state, as when their numbers are increased due to chronic stimulation, they become more dysfunctional. One possible culprit for chronic Th17 cell stimulation could be the changes in the composition of microbiota, or dysbiosis that is caused by shifting balances in beneficial or harmful bacteria species (Omenetti and Pizarro, 2015), as seen during HIV infection (Lujan et al., 2019), and as contributes to autoimmune diseases (Gulden et al., 2015). There is also evidence that Th17 cells can respond to specific microbiota-associated bacteria such as *Prevotella* species (Larsen, 2017). Indeed, several reports have identified microbiome changes in ME/CFS (Shukla et al., 2015;Giloteaux et al., 2016;Navaneetharaja et al., 2016;Nagy-Szakal et al., 2017;Proal and Marshall, 2018).

Regulatory T cells (Tregs) are tasked with suppressing autoimmune and excessive chronic inflammatory responses (Sakaguchi et al., 2010;Yamaguchi et al., 2011). Our finding that Tregs are increased in ME/CFS patients is consistent with our other findings that there appears to be a chronic activation of major T cell subsets with yet to be identified stimuli. Increased Tregs in patients may reflect the disturbance of the Th17 and Th1/Th2 cytokine balance in ME/CFS patients, suggesting a major disruption in homeostatic maintenance of immune responses. Indeed, for example, the balance between Th17 cells and Tregs is also critical in the regulation of inflammation, especially related to the gut and microbiome (Omenetti and Pizarro, 2015;Pandiyan et al., 2019). Remarkably, we found a correlation between Th17 and Treg cells in ME/CFS patients, and found the ratio was even more significantly different in patients compared to controls. These findings reveal potential biomarkers that can be utilized in future clinical interventions to re-balance the microbiome and the mucosal immune responses. In future studies it will also be important to further investigate in more detail the different subsets of Tregs for their suppressive capacity, and as with other subsets, to identify the mechanisms that lead to their increase in ME/CFS patients.

In our analysis, another T cell subset that was profoundly different between patients and controls, both phenotypically and functionally, was MAIT cells. MAIT cells selectively respond to a broad range of bacteria that possess the biosynthetic pathway for riboflavin metabolism (Gold et al., 2010;Le Bourhis et al., 2010;Kjer-Nielsen et al., 2012;Meierovics et al., 2013). A recent study found significant changes in the frequency of CD8+ MAIT cells in Multiple Sclerosis and severe ME/CFS patients (Cliff et al., 2019). We did not observe significant changes in the frequency of MAIT cells as a percentage of PBMC or T cells, which could be due to fact that this T cell subset is highly variable and has up to a 40-fold difference in range, even among healthy humans (Ben Youssef et al., 2018), and that most of our patient group was not characterized as severe. However, we observed profound differences in MAIT cell functions and differentiated states in ME/CFS subjects, such as reduced production of cytokines (IFNγ, IL-17) or cytotoxic molecule GranzymeA, and an increased proportion of CD27 negative CD8+ MAIT cells, which correlated with their lower survival in 6 day culture with IL-7 in vitro. Given that MAIT cells are specifically activated by a bacteria-produced riboflavin (vitamin B2) metabolite, we reason that this perturbation in MAIT cells can also be associated with differences in the composition of the microbiota of the patients. Indeed, we recently showed that MAIT cells can respond to a variety of microbiome-related bacteria (Tastan et al., 2018) and that they potentially function to tune or sense the microbial ecosystem throughout mucosal tissues. Two recent papers also show that MAIT cell development and expansion is also directly dependent on the microbiome (Constantinides et al., 2019;Legoux et al., 2019;Oh and Unutmaz, 2019). In one paper, the authors show that CD4-CD8- (double negative, or DN) MAIT cells are a functionally distinct subset that migrate to the skin in mice and are potentially involved in tissue repair (Dias et al., 2018;Constantinides et al., 2019;Oh and Unutmaz, 2019). Thus, it is interesting to note that we observed differences in CD8+ but not in DN MAIT cells in ME/CFS subjects, suggesting that these are different lineages or subsets. Taken together, it is conceivable that a disruption in the microbiome results in chronic activation of MAIT cells and an exhausted state in ME/CFS patients, which is similar to what has been seen with the Th17 subset (Cliff et al., 2019).

Finally, using immune parameters as features, our machine learning classifier was able to identify the ME/CFS patients at a high sensitivity and accuracy when using all 65 features, all 40 significantly different features and the 10 features among the 40 significantly different features that had the highest importance score. For all cases, we observed a higher value of sensitivity than specificity, indicating that the proportion of patients identified as ME/CFS patients is higher than healthy controls who are correctly identified as healthy. One reason for this could be related to features such as age, which causes the older individuals’ immune profiles to become more similar to those of ME/CFS patients, and hence the RF classifier categorizes healthy controls as patients. Currently, the diagnosis of ME/CFS is based on clinical symptoms alone and runs the potential for false positives and negatives. A classifier based on immune profiles might be an objective solution to better diagnose this clinical problem, but additional testing on larger and more diverse clinical cohorts will be required to assess the potential clinical utility of such an approach.

In conclusion, our findings open an exciting path to develop a set of biomarkers that can be utilized to aid in both diagnosis and in stratifying the patient population for targeted or precision medicine therapeutics.

## Data Availability

The source flow cytometry data for each subject underlying Figs. 1–8 and supplemental figures and tables will be provided upon reasonable request.

## Ethics Statement

This study was carried out in accordance with the recommendations of the Jackson Laboratory human subject ethics committee, and IRB protocol approval by Western Institutional Review board. All subjects gave written informed consent in accordance with the Declaration of Helsinki.

## Author Contribution

E.K. and C.L.G. contributed equally. E.K. processed blood samples, performed the majority of the experiments and analysis, and helped with the preparation of the manuscript; C.L.G. performed the statistical analyses and helped with the preparation of the manuscript; V.R. and J.G. performed machine learning analysis; P.N.G provided advice and feedback for machine learning analysis and interpretation; S.R., M.H. and L.P helped with processing blood samples, performed experiments and analysis; L.K., designed and quality checked all the immune panels; L.B. and S.D.V. recruited all subjects and clinically diagnosed ME/CFS patients; D.U. conceived and supervised the study and wrote the manuscript. All authors reviewed and approved the final draft of the manuscript.

## Acknowledgements

The research in this study was supported by National Institute of Health (NIH) grants R01AI121920 to D.U and U54 NS105539 to D.U, S.D.V., L.B and P.R.

## Abbreviations

ME/CFS: myalgic encephalomyelitis/chronic fatigue syndrome
PBMC: peripheral blood mononuclear cells
DN: double negative; CD4-CD8-
NK: natural killer
MAIT: mucosal-associated invariant T
Treg: T regulatory cells
N: naive
CM: central memory
EM: effector memory
EMRA: effector memory RA
PMA: phorbol 12-myristate 13-acetate
p: probability value
*r*_*s*_: Spearman’s Rank Correlation Coefficient
K: number of folds in cross validation
k: number of features
RF: random forest
TPR: true positive rate
FPR: false positive rate
ROC: receiver operating characteristic
AUC: area under the curve
F1: F score; measure of a test’s accuracy
EBV: Epstein-Barr virus
CMV: cytomegalovirus
HIV: human immunodeficiency virus

---

**Supplemental Figure 1.** (a) The frequency of each subset was correlated to subject age for CD4+ T cells, with clinical groups compared by nonparametric Spearman correlation, with exact *r*_*s*_ and p-value shown. (b) Analysis of CD4+ T cell subset frequencies in healthy control and ME/CFS patients that have been divided into two groups based on ages older and younger than 50 years. Data from healthy controls (Healthy, n = 91) and ME/CFS patients (ME/CFS, n = 190) for a and b, and clinical groups were compared by Mann-Whitney test for non-parametric data, with exact p values shown.

