## Supplementary figures and images for "Perturbation of effector and regulatory T cell subsets in Myalgic Encephalomyelitis/Chronic Fatigue Syndrome (ME/CFS)"

### Supplemental Figure 1

#### Supplemental figure 1

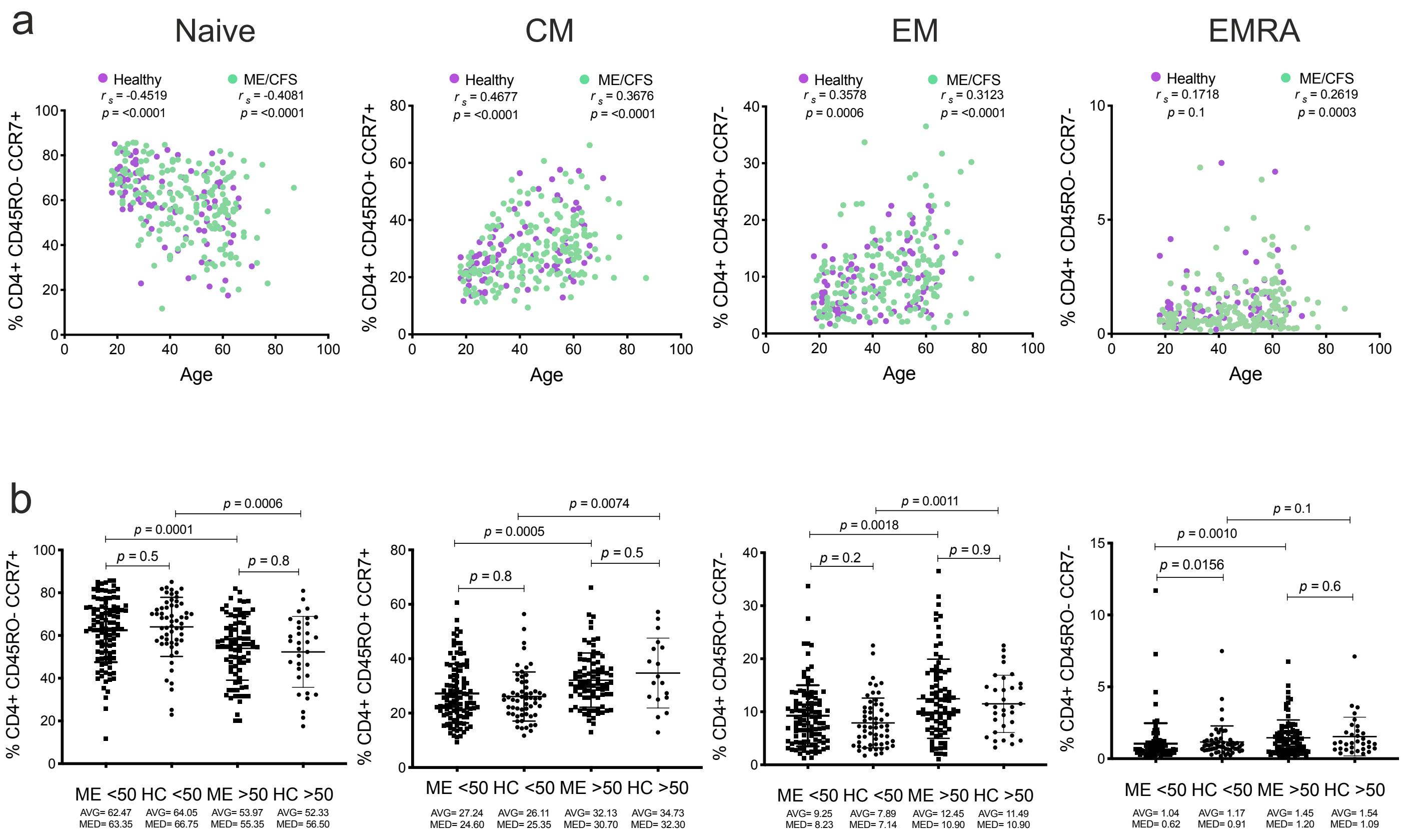
